# Intranasal immunization with a vaccinia virus vaccine vector expressing pre-fusion stabilized SARS-CoV-2 spike fully protected mice against lethal challenge with the heavily mutated mouse-adapted SARS2-N501Y_MA30_ strain of SARS-CoV-2

**DOI:** 10.1101/2021.12.06.471483

**Authors:** Karen V. Kibler, Mateusz Szczerba, Douglas Lake, Alexa J. Roeder, Masmudur Rahman, Brenda G. Hogue, Lok-Yin Roy Wong, Stanley Perlman, Yize Li, Bertram L. Jacobs

## Abstract

The Omicron SARS-CoV-2 variant has been designated a variant of concern because its spike protein is heavily mutated. In particular, Omicron spike is mutated at 5 positions (K417, N440, E484, Q493 and N501) that have been associated with escape from neutralizing antibodies induced by either infection with or immunization against the early Washington strain of SARS-CoV-2. The mouse-adapted strain of SARS-CoV-2, SARS2-N501Y_MA30_, contains a spike that is also heavily mutated, with mutations at 4 of the 5 positions in Omicron spike associated with neutralizing antibody escape (K417, E484, Q493 and N501). In this manuscript we show that intranasal immunization with a pre-fusion stabilized Washington strain spike, expressed from a highly attenuated, replication-competent vaccinia virus construct, NYVAC-KC, fully protected mice against disease and death from SARS2-N501Y_MA30_. Similarly, immunization by scarification on the skin fully protected against death, but not from mild disease. This data demonstrates that Washington strain spike, when expressed from a highly attenuated, replication-competent poxvirus, administered without parenteral injection can fully protect against the heavily mutated mouse-adapted SARS2-N501Y_MA30_.

## Introduction

The recently identified Omicron variant of SARS-CoV-2 has been designated a variant of concern because of its highly mutated spike protein (1). Of particular concern, Omicron spike is mutated at 5 positions (K417, N440, E484, Q493 and N501) that have been associated with escape from neutralizing antibodies induced by either infection with or immunization against the early Washington strain of SARS-CoV-2 (see Table 1)(2–4). Thus, Omicron may be able to at least partially escape from immunization with the current vaccines, which are all based on early, unmutated spike proteins.

**Table 1.** RBD Mutations.

| Table 1. RBD Mutations |  |  |  |  |  |
| --- | --- | --- | --- | --- | --- |
|  | Beta | Gamma | Delta | Omicron | SARS2-N501Y<sub>MA30</sub> |
| G339 | - | - | - | G339D | - |
| S371 | - | - | - | S371L | - |
| S373 | - | - | - | S373P | - |
| S375 | - | - | - | S375F | - |
| K417* | K417N | K417T | - | K417N | K417M |
| N440* | - | - | - | N440K | - |
| G446 | - | - | - | G446S | - |
| L452 | - | - | L452R | - | - |
| S477 | - | - | - | S477N | - |
| T478 | - | - | T478K | T478K | - |
| E484* | E484K | E484K | - | E484A | E484K |
| Q493* | - | - | - | Q493K | Q493R |
| G496 | - | - | - | G496S | - |
| Q498 | - | - | - | Q498R | Q498R |
| N501* | N501Y | N501Y | - | N501Y | N501Y |
| Y505 | - | - | - | Y505H | - |

\*Mutations associated with antibody escape.
[see (2-4)]

While the vaccines currently licensed or authorized for emergency use in the United States provide excellent protection against early variants of SARS-CoV-2, including Delta, they have limitations that may hinder their widespread worldwide use. They require maintenance of a significant cold-chain, and are administered parenterally, both of which may make widespread use difficult. We have generated a highly attenuated, replication-competent vaccinia virus vector, NYVAC-KC (5), which does not require an extensive cold-chain and can be administered either by scarification on the skin or intranasally (this manuscript). NYVAC-KC is fully replication competent in human primary keratinocytes and primary human dermal fibroblasts (5). Despite being replication competent, NYVAC-KC is highly attenuated in the very sensitive newborn intra-cranial mouse model, as well as in immune-deficient mice (5). NYVAC-KC induced mild induration on the skin of rabbits, with no signs of systemic spread (5). NYVAC-KC was highly immunogenic, inducing improved T cell and antibody responses to HIV inserts, compared to its replication deficient parental vector, NYVAC (5–10). Thus, NYVAC-KC may have properties that will make it useful in the worldwide fight against SARS-CoV-2.

In this manuscript we describe protection against challenge with a mouse-adapted variant of SARS-CoV-2, SARS2-N501Y_MA30_ (11). Early strains of SARS-CoV-2 are not pathogenic in mice. SARS2-N501Y_MA30_ was generated by serially passaging through mice of Washington strain SARS-CoV-2 that had an N501Y spike mutation. After 30 passages the virus became pathogenic for mice, which was associated with increased affinity for mouse ACE2 protein (11). During passage through mice 4 mutations accumulated in spike (along with 3 mutations in orf1a and 1 non-coding mutation in TRS), K417, E484, Q493, Q498 along with maintenance of the previous mutation at N501 (Figure 1). All 5 spike sites mutated in SARS2-N501Y_MA30_ are also mutated in Omicron, and 4 of the 5 mutated sites are at residues which when mutated allow escape from neutralizing antibodies induced by spike from early strains of SARS-CoV-2 (2–4). Thus, SARS2-N501Y_MA30_ expresses a highly mutated spike, which may also allow for escape from neutralizing antibodies induced by the current vaccines. However, we show that intranasal immunization with a pre-fusion stabilized Washington strain spike, expressed from the highly attenuated, replication-competent vaccinia virus vector NYVAC-KC, fully protected mice against both death and disease after infection with SARS2-N501Y_MA30_.Immunization by scarification fully protected against death, but not from mild disease. Thus, Washington strain spike, when expressed from a highly attenuated, replication-competent heat-stable poxvirus vector, administered without parenteral injection, can fully protect against challenge with the heavily mutated, mouse-adapted SARS2-N501Y_MA30_ variant of SARS-CoV-2.

**Figure 1.**
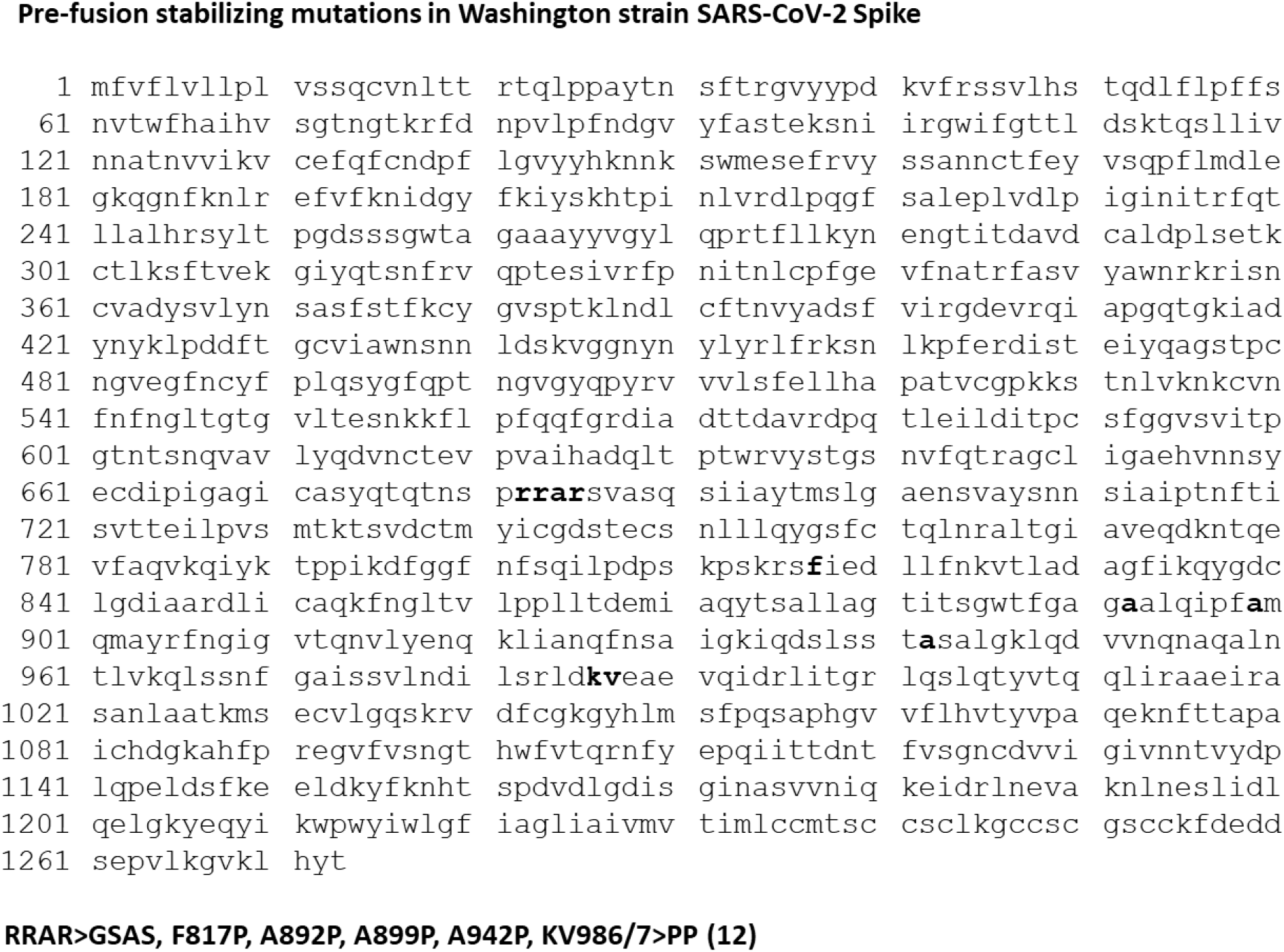
Pre-fusion stabilizing mutations in Washington strain Spike. The indicated mutations (in **bold**) were made to stabilize Washington strain Spike in the pre-fusion conformation, as previously described (12).

## Results

### Generation of NYVAC-KC-pfsSpike

A vaccinia virus-optimized Washington strain spike was stabilized in the pre-fusion state by mutation of the furin cleavage site, and insertion of 6 proline residues, preventing the conformational change to the post-fusion conformation (pfsSpike) (12) (Figure 1). PfsSpike, flanked by TK locus homologous flanking arms, was inserted into the TK locus of NYVAC-KC by homologous recombination (Figure 2). The TK locus of NYVAC-KC was modified by insertion of a pGNR-cmr^S^ cassette (13) prior to homologous recombination with TK flanked pfsSpike. pGNR-cmr^S^ encodes a neo^r^ gene and a GFP gene, to allow for selection and identification of virus that has taken up pGNR-cmr^S^, as well as a cmr^S^ gene that acts as a negative selectable marker (14). Cells were infected with NYVAC-KC-neo^R^-GFP-cmr^S^ and transfected with TK-flanked pfsSpike. Recombinant virus that had replaced the pGNR-cmr^S^ cassette with pfsSpike was selected for as cmr^R^, non-fluorescent plaques. Insertion of pfsSpike was confirmed by PCR and Western blot of individual plaques. This technology allows for rapid insertion (approximately 1 month from obtaining DNA to having a P2 stock) of new genes into NYVAC-KC.

**Figure 2.**
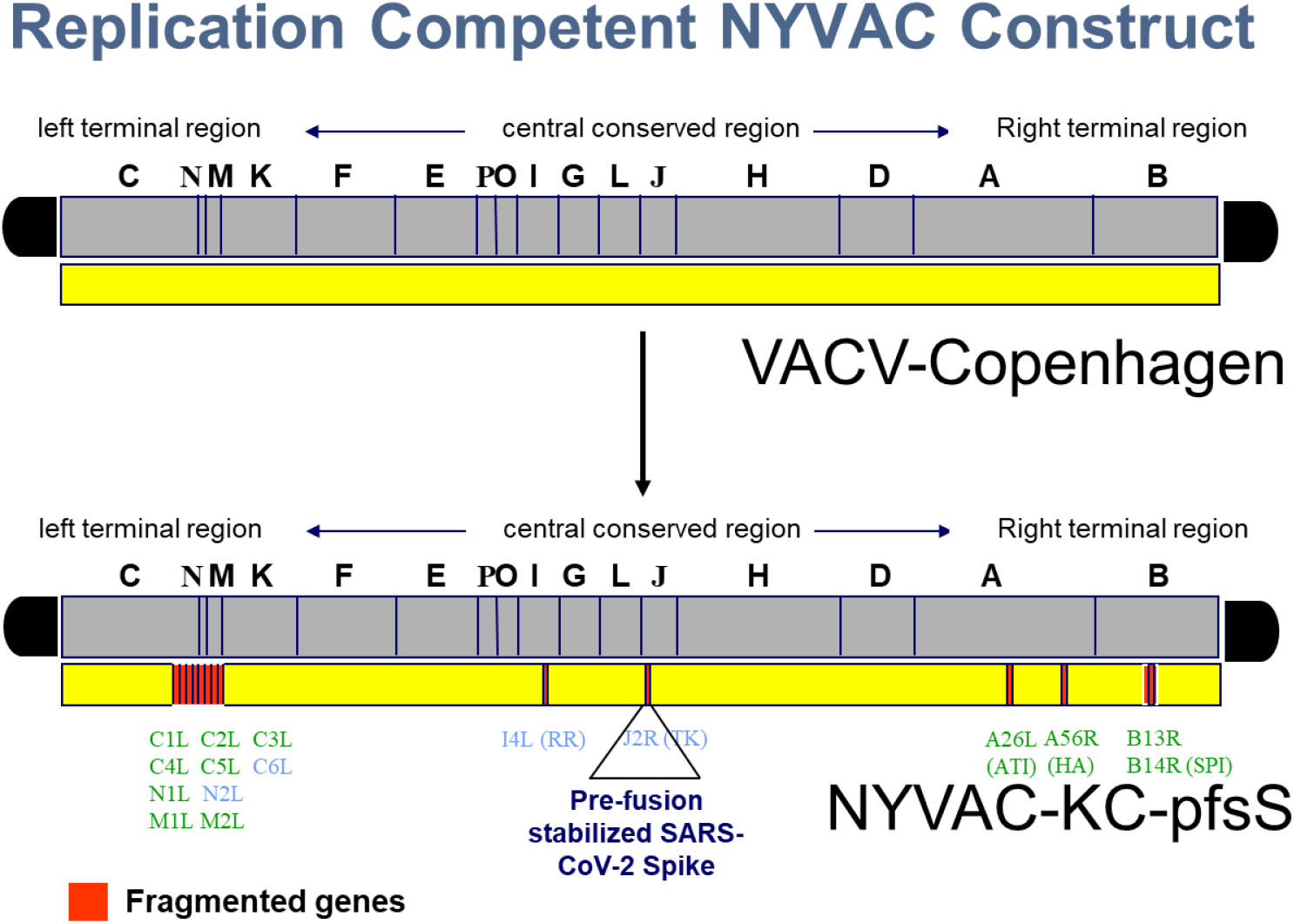
NYVAC-KC-pfsSpike. NYVAC-KC is a highly attenuated, replication-competent derivative of the Copenhagen strain of vaccinia virus, that has been deleted of 16 open reading frames. A pre-fusion stabilized spike, under control of a synthetic early/late promoter was inserted into the TK locus of NYVAC-KC to generate NYVAC-KC-pfsSpike.

### Immunization with NYVAC-KC

Mice were immunized with 10^6^ pfu of NYVAC-KC-pfsSpike, either by scarification or intranasally (Figure 3). Mice were boosted at one month post immunization, rested for 3 months, and boosted a second time. Blood was obtained one month after the primary immunization, one and three months after the first boost and two weeks after the second boost. Serum was assayed for the ability to block binding of Washington strain Spike protein RBD to human ACE2 (15). Immunization by scarification with NYVAC-KC-pfsSpike gave a modest serum response inhibiting RDB binding to huACE2 (Figure 4A). The response was boosted to high levels, which waned after three months. The second boost increased the serum response, inhibiting binding of RBD to huACE to moderate levels. A single intranasal immunization with NYVAC-KC-pfsSpike induced a potent serum response that inhibited RDB binding to huACE2 (Figure 4B). This response remained high after the first boost and did not appreciably wane three months after the first boost, and remained high after the second boost. Thus, intranasal immunization was able to induce a potent durable serum RBD binding response.

**Figure 3.**
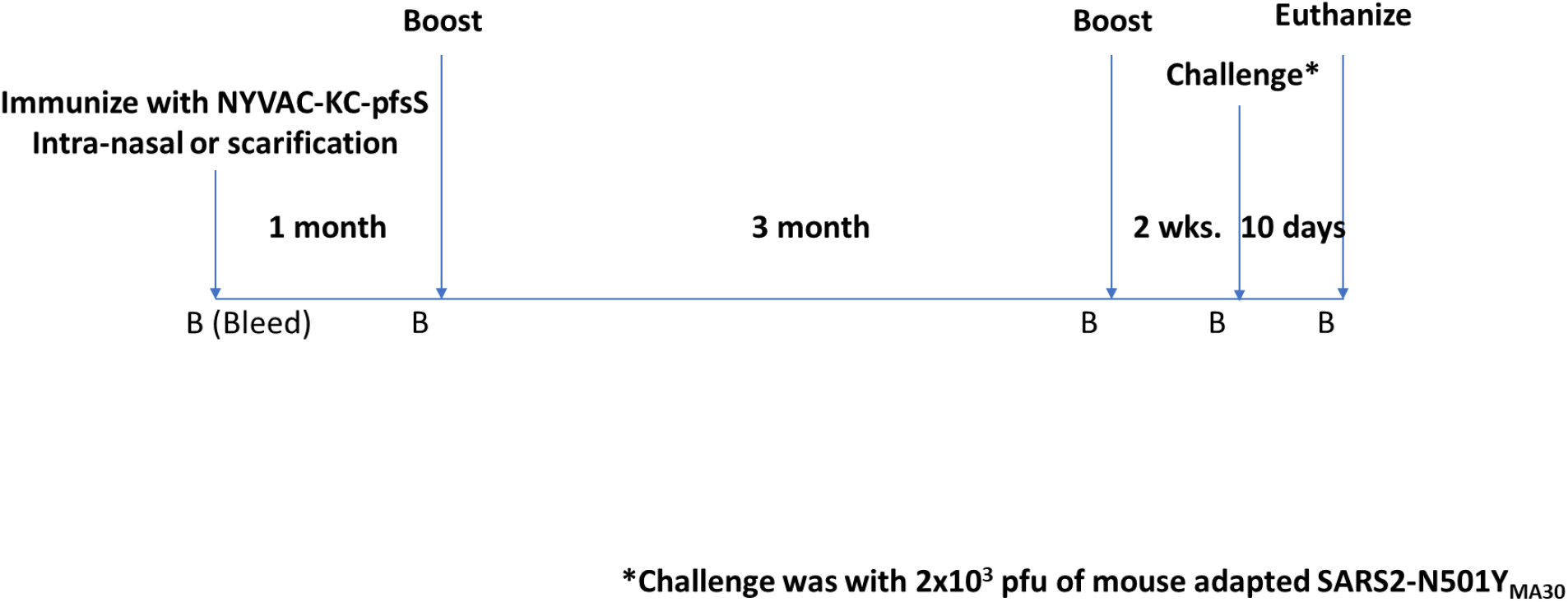
Immunization/challenge schedule. Animals were immunized on day 0 and boosted 1 month and 4 months after the first immunization. Animals were challenged 2 weeks after the second boost and monitored for signs of morbidity for up to 10 days. Animals were bled (indicated by “B”) one day prior to each immunization, one day prior to challenge and for all surviving animals, at the termination of the experiment.

**Figure 4.**
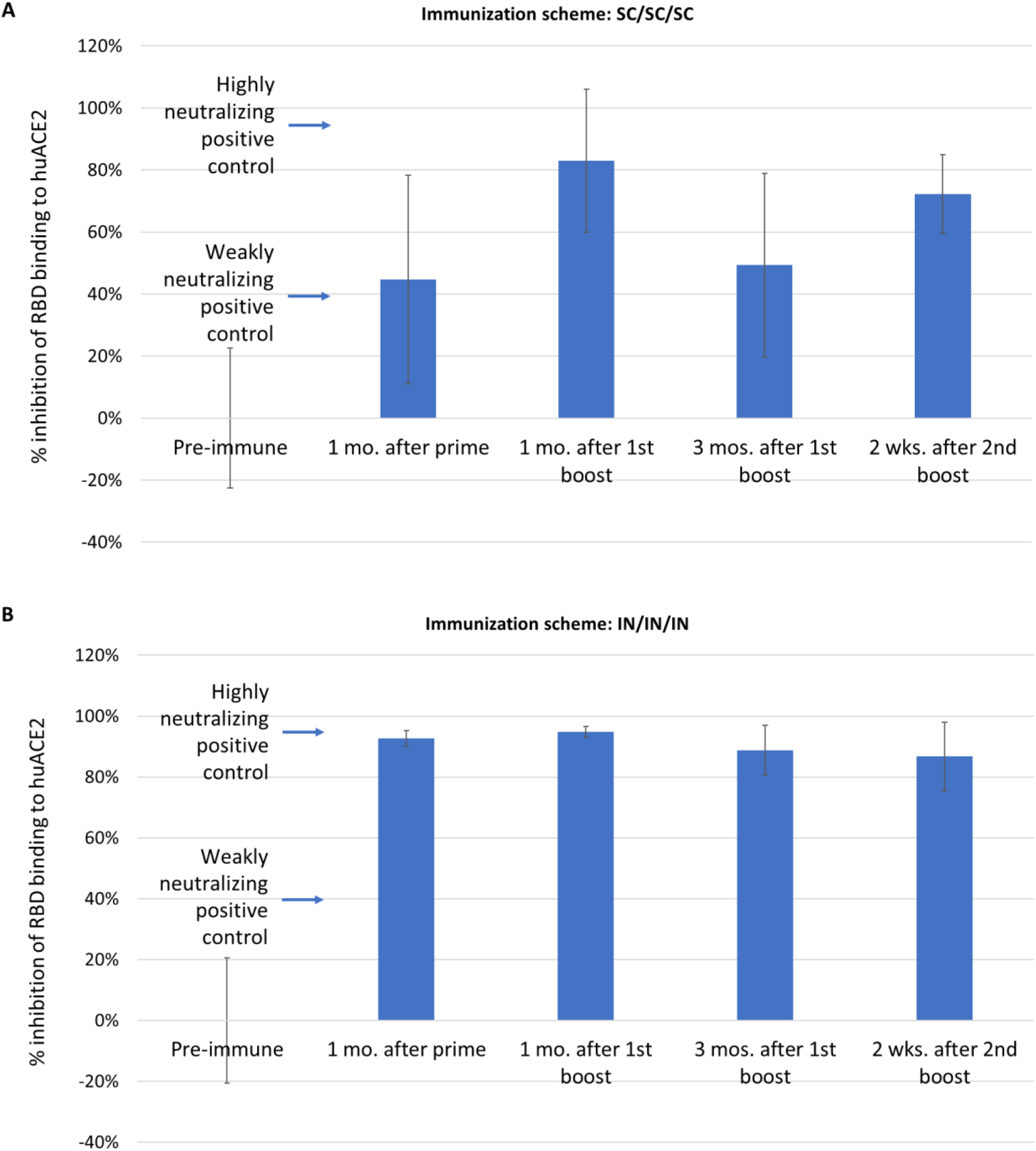
RBD binding antibodies. Serum from animals bled at the indicated times were assayed for the ability to inhibit binding of gold-labeled Washington strain RBD to huACE2. Controls indicated inhibition of binding by a strongly neutralizing positive control, and a weakly neutralizing positive control.

### Challenge with SARS2-N501Y_MA30_

Animals immunized with NYVAC-KC-pfsSpike were challenged two weeks after the second boost with approximately 2×10^3^ pfu of SARS2-N501Y_MA30_ (11). Animals were monitored and scored from 0-3 according to severity for each criterion: weight loss, ruffled fur, hunching, and loss of activity. All animals were scored in a blinded fashion. An aggregate clinical score of 8 was an endpoint for humane euthanasia. Fifteen of seventeen animals not immunized with NYVAC-KC-pfsSpike reached a clinical score of 8 by 4 days post-infection and were humanely euthanized (Figure 5, red line). None of the animals immunized with NYVAC-KC-pfs-Spike needed to be euthanized (Figure 5, blue line). Figure 6 shows the clinical score for each animal in aggregate groups from 0-9 days post challenge. Mock challenged animals had scores of 0-1 throughout the course of the experiment (Figure 6A). Animals not immunized with NYVAC-KC-pfsSpike, and challenged with SARS2-N501Y_MA30_, all showed signs of illness by days 2-3 post-challenge, and for 15 of the 17 animals, symptoms were serious enough to warrant humane euthanasia (Figure 6B). Animals immunized with NYVAC-KC-pfsSpike were all spared progression to serious disease (Figure 6C), although two of the animals had mild disease, with maximal clinical scores of 2 and 4. The two animals that showed mild disease were in the cohort that was immunized by scarification (Figure 6D). Intranasally immunized animals were asymptomatic after SARS2-N501Y_MA30_challenge, with clinical scores of 0-1 (Figure 6E), indistinguishable from mock challenged animals (Figure 6A).

**Figure 5.**
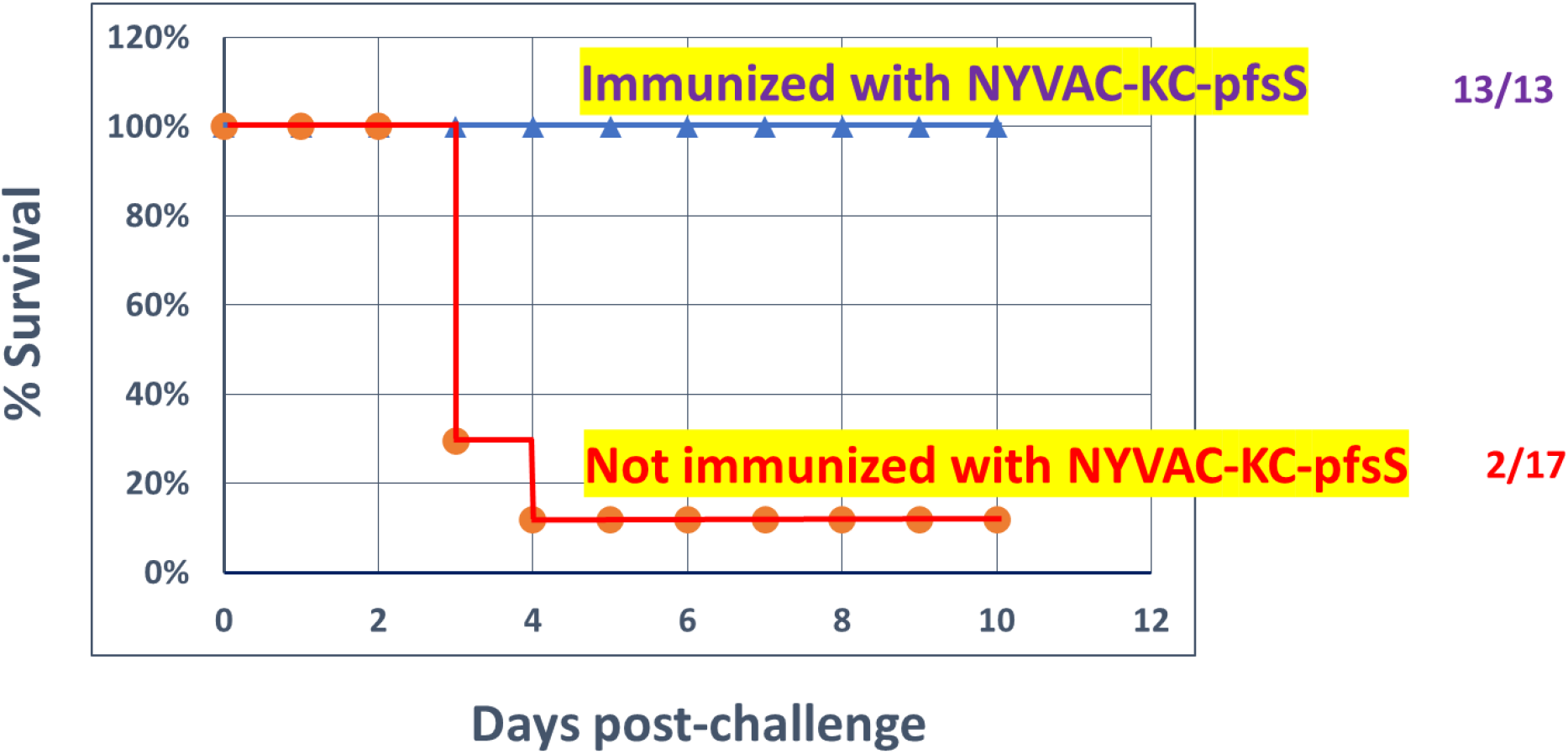
Survival after challenge with mouse-adapted SARS-CoV-2, SARS2-N501Y_MA30_. Animals either not immunized with NYVAC-KC-pfsSpike or immunized with NYVAC-KC-pfsSpike were challenged with 2×10^3^ pfu of mouse adapted SARS2-N501Y_MA30_. Animals were monitored for morbidity daily in a blinded manner for up to 10 days (see Figure 6). Animals with a clinical score of 8 or higher were humanely euthanized.

**Figure 6.**
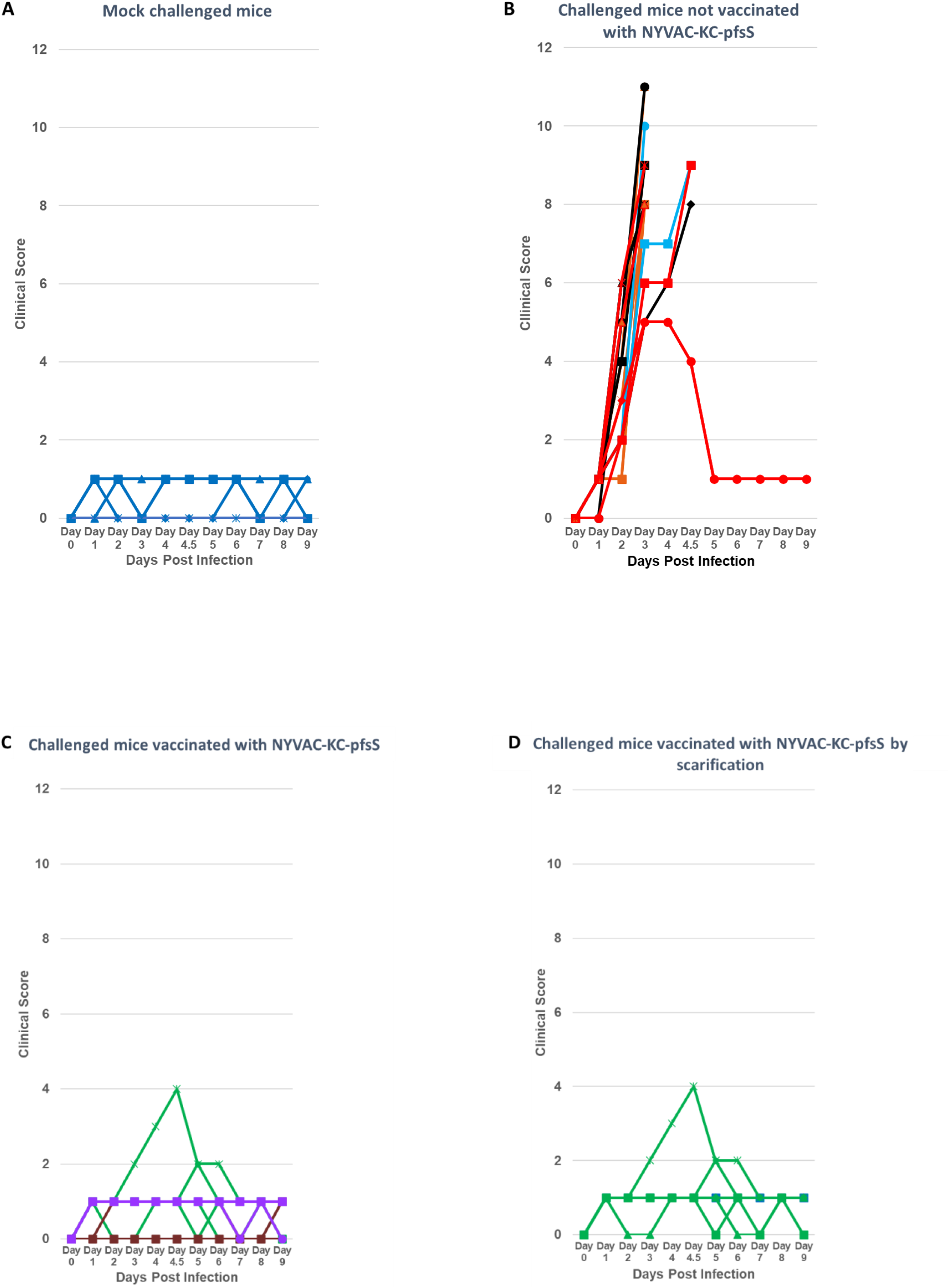
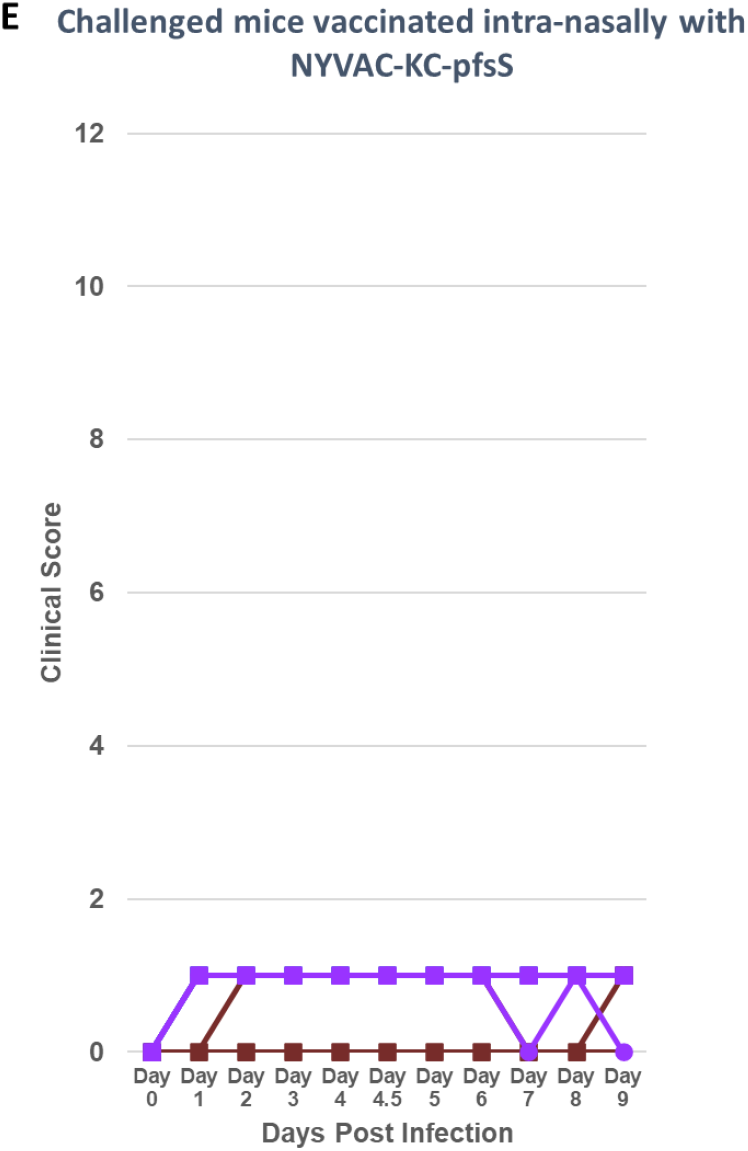
Clinical scores of challenged animals. Animals were monitored for morbidity (weight loss, ruffled fur, hunching, diminished activity, with a range or 0-3 for each parameter, with 0 being no symptoms, 1, mild symptoms, 2, moderate symptoms, and 3, severe symptoms) for up to 10 days after challenge. Animals with an aggregate score of 8 or greater were humanely euthanized. **A.** Animals not immunized with NYVAC-KC-pfsSpike and not challenged. **B.** Animals not immunized with NYVAC-KC-pfsSpike and challenged with mouse adapted SARS2-N501Y_MA30_. **C.** Animals immunized with NYVAC-KC-pfsSpike and challenged with mouse adapted SARS2-N501Y_MA30_. **D.** Animals immunized by scarification with NYVAC-KC-pfsSpike and challenged with mouse adapted SARS2-N501Y_MA30_. **E.** Animals immunized intranasally with NYVAC-KC-pfsSpike and challenged with mouse adapted SARS2-N501Y_MA30_.

## Discussion

In this manuscript we demonstrate that Washington strain spike, expressed from a highly attenuated, replication-competent vaccinia virus vector, NYVAC-KC, fully protected mice from serious disease after challenge with a heavily mutated, mouse adapted strain of SARS-CoV-2, SARS2-N501Y_MA30_. NYVAC-KC was originally developed as an improved poxvirus-based vector for immunization against HIV (5). NYVAC-KC is highly attenuated yet induces a potent T cell and antibody response against HIV *gag, pol, nef* and *env* inserts (5–10). Since poxvirus vectors are heat stable and generally don’t require an extensive cold-chain (16), NYVAC-KC based vectors likely will be easy to distribute worldwide. In this manuscript we demonstrate that NYVAC-KC expressing a pre-fusion stabilized spike can induce protective immune responses when administered either by scarification or intranasally. Thus, these vectors may be easy to administer after widespread distribution. Furthermore, multiple antigens can be expressed from NYVAC-KC. There are 6 deletion sites in NYVAC-KC (Figure 2), each of which can be used to express foreign antigens. It is also possible to express multiple antigens from each insertion site. We have successfully generated a stable construct expressing HIV *gag*, *pol*, *nef* and *env* from the TK locus (5). Thus, it may be possible to express spikes from multiple SARS-CoV-2 variants in a single vector, and to express antigens encoding stable T cell epitopes, in addition to the highly variant Spike proteins.

Early strains of SARS-CoV-2 do not cause disease in wild type mice. On the contrary, mouse adapted SARS2-N501Y_MA30_ is highly virulent in wild type mice (11). Intranasal administration of a dose of 2×10^3^ pfu uniformly induced serious disease in infected animals, with the majority of animals being euthanized by 4 days post-infection. During the course of adaptation in mice, SARS2-N501Y_MA30_ fixed 5 mutations in the Spike RBD (11). These mutations were associated with increased binding to mouse ACE2. Interestingly, while SARS2-N501Y_MA30_ was selected for in immunologically naïve mice, 4 of the 5 RBD mutations are at loci associated with resistance to neutralizing antibodies induced by early strain Spike (2–4). Thus, these mutations may have multiple effects, enhancing binding to murine ACE2, while possibly providing at least partial resistance to neutralizing antibodies. All 5 of the loci in RBD with fixed mutations in SARS2-N501Y_MA30_ are also mutated in the heavily mutated Omicron variant of SARS-CoV-2 (Table 1).

Immunization with NYVAC-KC-pfsSpike fully protected mice from lethal challenge with the heavily mutated SARS2-N501Y_MA30_, despite the immunogen in NYVAC-KC-pfsSpike having a wild type, unmutated RBD. Thus, either NYVAC-KC-pfsSpike induces neutralizing antibodies to regions not mutated in SARS2-N501Y_MA30_ spike, or induces a high enough neutralizing antibody titer to cross-neutralize the divergent SARS2-N501Y_MA30_. While we are in the process of measuring neutralizing antibody levels induced by NYVAC-KC-pfsSpike, we have shown that NYVAC-KC-pfsSpike can potently induce antibodies that inhibit binding of Washington strain RBD to human ACE2 (Figures 4A and B). It is not clear if these antibodies can inhibit binding of SARS2-N501Y_MA30_ RBD to mouse ACE2, or if NYVAC-KC-pfsSpike induces antibodies to other regions of Spike.

While either intranasal or scarification immunization with NYVAC-KC-pfsSpike protected mice from serious disease induced by SARS2-N501Y_MA30_,intranasal immunization appeared to give superior protection, with animals being fully asymptomatic after challenge. Intranasal immunization in animals also induced more potent serum antibody responses that inhibited binding of Washington strain SARS-CoV-2 Spike RBD to human ACE2, than immunization by scarification. It is unclear if these higher titers of serum RBD binding antibodies led to enhanced protection after intranasal immunization, or if intranasal immunization led to an enhanced mucosal immune response that fully protected against disease.

In conclusion, in this manuscript we demonstrate that pre-fusion stabilized Washington strain spike, when expressed from a highly attenuated, replication-competent poxvirus vector, administered without parenteral injection can fully protect against the heavily mutated mouse-adapted SARS-CoV-2, SARS2-N501Y_MA30_.

## Materials and Methods

### Viruses

Mouse adapted SARS-CoV-2 SARS2-N501Y_MA30_ was propagated in A549-huACE2 cells (11). For insertion of foreign genes into the NYVAC-KC genome, we constructed a cassette (pGNR-cmr^S^) that encodes an *E*. coli gyrase/PKR fusion protein that confers coumermycin (cmr) sensitivity (14), a neo^R^ gene and expresses GFP (13). The cassette has arms that are homologous to the sequence flanking the TK deletion in NYVAC-KC, to allow for in vivo recombination with the viral genome. The pGNR-cmr^S^ cassette was added to NYVAC-KC through an in vivo recombination (17) done in BSC-40 cells; cells were transfected with linear cassette DNA using Lipofectamine 2000 (Invitrogen) according to product instructions. Infection with NYVAC-KC was at an MOI of 0.05. After 48 hours, the infected cells were scraped into the medium (1.2 mls Opti-Pro (Gibco) with glutamine and 1% FBS). Following two cycles of freeze/thaw, the cell supernatant was used to infect 100 mm dishes of BSC-40 cells, at 1:10, 1:100, and 1:1000 dilutions of the IVR stock. DMEM 2% FBS plus G418 at 1 mg/ml was added after the infection incubation. Green, G418^R^ plaques were picked at 48 hours post infection, following the addition of an agarose overlay. Plaques were screened in 6-well plates for sensitivity to cmr, and the two showing the highest sensitivity were chosen for continuing to the next round of plaque purification in BSC-40 cells. The plaque from this round that demonstrated the highest sensitivity to cmr was amplified in a 60 mm dish. This virus (NYVAC-KC-neo^R^-GFP-cmr^S^) was used in an IVR to replace the pGNR-cmr^S^ cassette with the coding sequence for a vaccinia virus optimized, pre-fusion-stabilized SARS-CoV-2 Washington strain spike protein (12), under control of a vaccinia virus synthetic early/late promoter (18), yielding a cmr^R^, non-fluorescent virus. For this selection, 100 ng/ml cmr was added at 24 hpi of the IVR, and subsequent infections were carried out in the presence of cmr until the final plaque was chosen. Correct insertion was confirmed by PCR and Western blotting. Plaques were amplified twice to obtain P2 stocks (5) that were used for immunization of mice.

### Cell lines

African green monkey kidney Vero cells (E6) or (CCL81) (obtained from ATCC) were cultured in Dulbecco’s modified Eagle’s medium (DMEM; Gibco catalog no. 11965), supplemented with 10% fetal bovine serum (FBS), 100 U/ml of penicillin, 100 μg/ml streptomycin, 50 μg/ml gentamicin, 1mM sodium pyruvate, and 10mM HEPES. Human A549 cells (Verified by ATCC) were cultured in RPMI 1640 (Gibco catalog no. 11875) supplemented with 10% FBS, 100 U/ml of penicillin, and 100 μg/ml streptomycin. The generation of A549-ACE2 cells was described previously (19).

### Plaque assay

Briefly virus supernatant was serially diluted 10-fold and inoculum was absorbed on Vero cells for 1 hour at 37°C. Inoculum was overlaid with DMEM plus 0.7% agarose and incubated for 3 days at 37°C. Cells were fixed with 4% paraformaldehyde and stained with 1% crystal violet for counting plaques. All infections and virus manipulations were conducted in a biosafety level 3 (BSL-3) laboratory using appropriate and IBC-approved personal protective equipment and protocols.

### Immunization

BALB/c mice at age 7 weeks were immunized with 10^6^ pfu of NYVAC-KC-pfsSpike. Immunization was performed either intranasally (in 10 μL), or by tail scarification (20 μL) and under anesthesia with a cocktail containing 37.5 mg/kg ketamine, 7.5 mg/kg xylazine, and 2.5 mg/kg acepromazine. Following vaccination, mice were allowed to recover on heating pads and were monitored until ambulatory, at which point they were placed in their cages. Mice were boosted 1 month and 4 months after initial vaccination. Throughout the duration of the study before challenge, mice were weighed weekly and blood draws were taken on a bi-weekly basis.

### Inhibition of RBD/huACE2 interaction

Neutralizing antibodies were assessed using a lateral flow assay that semi-quantitatively measures levels of antibodies that prevent binding of Washington strain RBD to ACE2, as previously described (15). Briefly, 3 μl of serum was diluted to 6 μl in PBS and loaded onto lateral flow strips that had soluble gold-labeled Washington strain RBD, and bound huACE2. Serum and gold-labeled RBD were chased through the strip with chase buffer (15). After 20 minutes, blue color at the site of the bound ACE2 was quantified by densitometry. Percent inhibition was calculated as previously described (20), using the following formula: 1-(Test sample line density/Limit of Detection, LoD)*100 where LoD for non-neutralizing sera for the rapid test was 570,229.

### SARS2-N501Y_MA30_ Challenge

Mice either immunized or not immunized with NYVAC-KC-pfsSpike were moved to the ABSL3 for SARS-CoV-2 challenge. SARS2-N501Y_MA30_ was administered intranasally at a dose of 2×10^3^ pfu per animal in a volume of 50 μl. Mice were anesthetized by intraperitoneal route with a cocktail of 50 mg/kg ketamine and 7.5 mg/kg xylazine for the inoculation. Following the inoculation, mice were allowed to recover in their cages, which were placed on heating pads, and mice were monitored until ambulatory. Mice were weighed daily unless weight fell below 85% of their original weight, at which time they were monitored twice daily. Symptoms were scored in a blinded manner for ruffled fur, hunching and activity, and scored from 0-3 (0 normal, 3 severe) for 10 days and mice were euthanized when their aggregate clinical score reached 8 (including a score of 0-3 for weight loss) as detailed in the approved IACUC protocol. Mice that recovered or were asymptomatic were monitored for 10 days.

